# PGRanker: Enhancing Phenotype Knowledge with Graph Learning for Mendelian Gene Prioritization

**DOI:** 10.64898/2026.03.11.711106

**Authors:** Ryan Köksal, Adrian Fritz, Miriam Schmidts, Anup Kumar, Van Dinh Tran, Rolf Backofen

**Affiliations:** Bioinformatics Lab, Albert-Ludwigs Universität Freiburg, Georges-Köhler-Allee 101, 79110, Freiburg im Breisgau, Germany; Center for Pediatrics and Adolescent Medicine, University of Freiburg Medical Center, Mathildenstr. 1, 79106, Freiburg im Breisgau, Germany; Research and Innovation, King Faisal Specialist Hospital & Research Centre, Riyadh, Saudi Arabia; Signalling Research Centre CIBSS, University of Freiburg, Schaenzlestr. 18, 79104, Freiburg, Germany

**Keywords:** phenotype, Mendelian, machine learning, disease gene prioritization, graph

## Abstract

**Motivation:** Phenotype-driven causal-gene prioritization depends on both the coverage of curated annotations and the quality of patient phenotype descriptions. Direct phenotype matching provides evidence for annotated genes, while gene networks can extend this evidence to functionally related candidates. Integrating these sources effectively, while maintaining performance under incomplete or imprecise phenotyping, remains an important challenge in Mendelian disease diagnosis.

**Results:** We present PGRanker, a phenotype-driven method that uses direct HPO–gene support to seed patient-specific graph learning. A residual graph-attention network refines this support through gene interactions, and a learned scorer combines the resulting gene representations with an attention-pooled phenotype profile to rank 21,592 candidate genes without variant data. We evaluated PGRanker against Phen2Gene, CADA and GADO on 277 diagnosed cases and a separate clinical cohort of 21 cases with sparse phenotype descriptions. On the 277-case benchmark, PGRanker achieved the highest recall at every evaluated cutoff, increasing mean reciprocal rank from 0.142 for the strongest baseline, CADA, to 0.192, a 35% relative improvement. Median causal-gene rank improved from 99 to 73, and recall@1 from 9.0% to 14.0%. PGRanker also achieved the highest recall@10 area under the degradation trajectory across four controlled perturbations: phenotype removal, addition of unrelated terms, substitution with unrelated terms, and replacement by less-specific ancestors. These results support patient-specific integration of curated phenotype evidence and gene-network learning for more accurate causal-gene prioritization across the tested phenotype-input conditions.

**Availability and implementation:** Code and data are available at https://github.com/ivelet/PGRanker.

## Introduction

Rare Mendelian disorders are individually uncommon but collectively affect a large population, and molecular diagnosis remains slow and incomplete despite major advances in genome sequencing. Large clinical studies continue to report diagnostic yields well below 50% [20, 6, 16], indicating that sequencing alone does not resolve many rare-disease cases. A central challenge is therefore no longer the generation of genomic data, but the prioritization of candidate genes that plausibly explain a patient’s clinical phenotype. Exome and genome sequencing can identify many rare and potentially damaging variants, while only a small fraction are relevant to the underlying disorder [3, 4]. Phenotypic similarity provides an important source of evidence because disorders with similar clinical manifestations are often associated with functionally related genes [19, 9].

The Human Phenotype Ontology (HPO) provides a standardized representation of phenotypic abnormalities and enables patient findings to be encoded as a set of computable terms [24, 15]. Together with curated disease and gene annotations in resources such as OMIM [1], HPO has enabled phenotype-driven gene prioritization, in which a patient’s observed phenotype profile is used to rank candidate genes. A major class of methods relies directly on curated phenotype-gene or phenotype-disease associations. Phenotype-driven prioritizers fall broadly into annotation-based and network-based approaches, with overlapping mechanisms and complementary strengths. Annotation-based approaches rank genes using curated term-to-gene associations or phenotype similarity measures. Early methods compared patient HPO terms with curated disease annotations using ontology-based semantic similarity [14]. Subsequent approaches weighted rare, specific terms more heavily than common ones [11], while likelihood-ratio frameworks scored diseases according to how well they explained the observed phenotypes, optionally incorporating variant evidence [25]. Other methods widen the evidence base: AMELIE mines primary literature to score genes against a patient’s phenotype [2], and Phen2Gene aggregates weighted HPO-term-to-gene associations from a precomputed knowledge base (H2GKB) without requiring variant information [29]. Direct lookup is computationally efficient and can be traced to the underlying associations. However, these approaches depend on the coverage of existing phenotype annotations. Causal genes with sparse or absent annotations may therefore receive little support, even when the patient’s phenotype is informative.

A complementary class of approaches exploits relationships among genes and other biological entities to extend phenotype evidence beyond directly annotated candidates. Network-propagation methods spread disease or phenotype information through molecular interaction networks [13, 5, 8], while graph neural networks learn representations and propagation mechanisms from biological graphs [17, 27, 18]. GADO predicts gene–phenotype associations using co-regulation with genes annotated to the relevant phenotypes [7], whereas CADA builds a case-enriched HPO–gene knowledge graph and performs prioritization through link prediction over learned node embeddings [21]. These approaches can provide evidence for candidates with limited direct phenotype annotation. Integrating this broader network information while retaining informative, patient-specific support from curated phenotype– gene associations is therefore a natural objective for gene prioritization.

Curated phenotype associations and gene networks provide complementary sources of evidence: the former connect observed phenotypes to characterized genes, while the latter can extend that evidence to functionally related candidates. This motivates an integration strategy in which patient-specific phenotype support determines the input to learned graph propagation. Such a model can retain evidence at directly supported genes while incorporating information from their interaction partners. Whether this refinement improves causal-gene ranking beyond direct lookup, and when it instead weakens informative predictions, requires explicit evaluation.

The usefulness of these approaches also depends on the quality of the supplied phenotype profile. Clinical descriptions may be incomplete, evolve over time, or be inconsistently translated into structured phenotype terms [10]. Evaluating gene prioritization therefore requires testing sensitivity to missing observations, incorrect terms, and reduced phenotypic specificity.

Here we present PGRanker, a phenotype-driven gene prioritizer that integrates direct HPO–gene support with patient-specific gene-network learning. For each patient, the fraction of supplied, indexed HPO terms associated with each candidate gene is projected into a node representation and propagated through a single residual graph-attention layer over HumanNet, with interaction confidences informing attention. This produces gene representations that combine each gene’s own phenotype support with evidence from its network neighbors. An attention-pooled representation of the patient’s HPO terms conditions a learned scorer that ranks all 21,592 candidate genes. Direct lookup thus supplies the evidence propagated through the network, making gene representations specific to the patient before scoring.

We evaluated PGRanker against Phen2Gene, CADA and GADO on Benchmark-277, comprising 277 solved cases from five independently ascertained cohorts. PGRanker achieved the highest recall at every evaluated cutoff, a mean reciprocal rank (MRR) of 0.192 and a median causal-gene rank of 73, compared with 0.142 and 99 for CADA, the strongest baseline. Recall@1 was 14.0%, compared with 9.0% for CADA. Comparisons with direct lookup examined the contribution and limitations of learned refinement, while a separate clinical cohort of 21 cases assessed performance across diverse monogenic disorders with sparse phenotype descriptions. Finally, we assessed sensitivity to phenotype degradation through HPO-term removal, addition of unrelated terms, substitution with unrelated terms, and replacement by less-specific ancestors. PGRanker achieved the highest area under the recall@10 trajectory in all four regimes.

## Methods

### Problem formulation

Given the HPO terms observed in a clinical case, PGRanker ranks a fixed genome-wide candidate set to prioritize the causal gene, without variant data or a pre-filtered candidate list.

Let *G* = *{g*_1_, …, *g*_*N*_ *}* denote the candidate-gene universe (*N* = 21,592) and *ℋ* the vocabulary of 8,419 HPO terms annotated to OMIM disorders across the training, validation and test partitions. For each case *i*, the input is a deduplicated set of HPO terms *Q*_*i*_, of which only the indexed subset 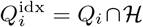 is used. Two external resources are held fixed: a gene–gene network with edges *E* and interaction confidences *w*, and a binary HPO–gene association matrix *A ∈ {*0, 1*}*^|*ℋ*|*×N*^ .

PGRanker learns a scoring function *s*_*θ*_(*g, Q*_*i*_; *A, E, w*) *∈* ℝ,where *θ* denotes the learned parameters and higher scores indicate greater predicted compatibility with the phenotype profile. For each case, the candidate ranking is

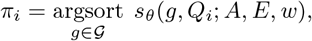

in descending score order, with ties resolved by ascending Entrez Gene identifier.

### PGRanker integrated pipeline

PGRanker integrates phenotype-gene evidence, gene-network learning, and phenotype-query encoding within a single ranking model (Figure 1). For each case, direct HPO–gene support is computed for every candidate gene and projected into node representations that initialize the graph-attention encoder over GeneNet, the gene network derived from HumanNet. A residual graph-attention layer combines each gene’s own support with evidence from its interaction partners. The network is fixed, but its inputs and resulting gene representations are specific to the case. In parallel, an attention-based query encoder pools the same indexed HPO terms into a phenotype-profile representation. A learned scorer combines this query representation with each gene’s graph-updated representation to rank all 21,592 candidate genes. Please refer to Supplementary Sections 1-3 for detailed descriptions of datasets and experimental setup, and baseline methods compared against.

**Figure 1.**
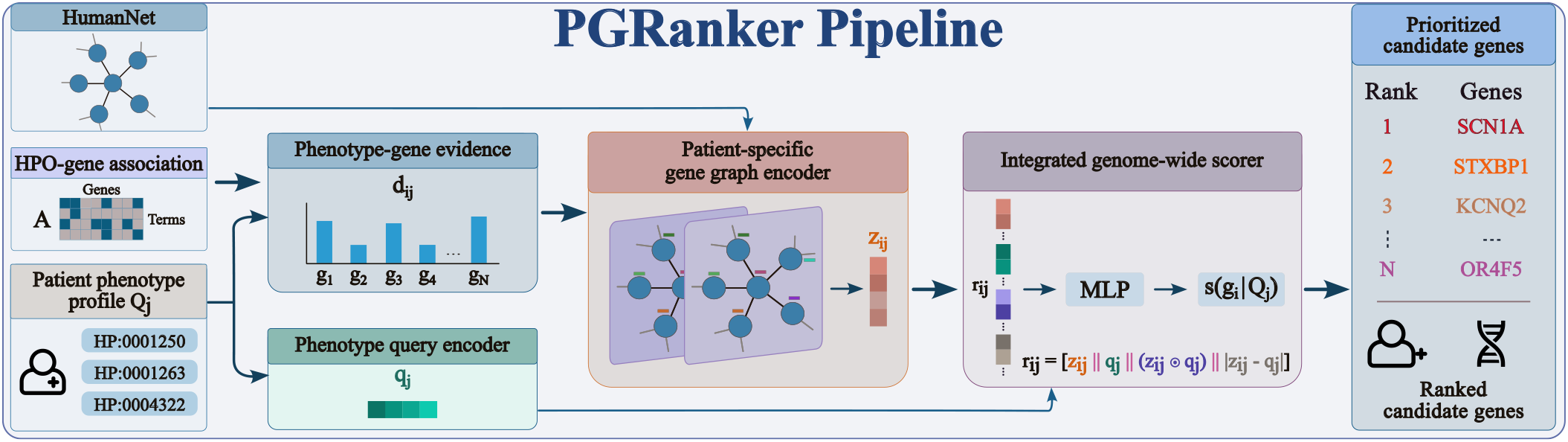
Overview of the PGRanker pipeline. For a patient phenotype profile *Q*_*j*_, a phenotype-gene evidence score *d*_*ij*_ is computed for each candidate gene *g*_*i*_ from the HPO-gene association matrix *A* and used as node input to a graph attention network over HumanNet, yielding a graph-refined representation **z**_*ij*_ . In parallel, the profile is encoded into a query representation **q**_*j*_ . The scorer concatenates **z**_*ij*_, **q**_*j*_, their element-wise product, and their absolute difference into **r**_*ij*_, and an MLP maps **r**_*ij*_ to a relevance score *s*_*θ*_(*g*_*i*_ | *Q*_*j*_ ); all 21,592 candidate genes are ranked by this score.

**Figure 2.**
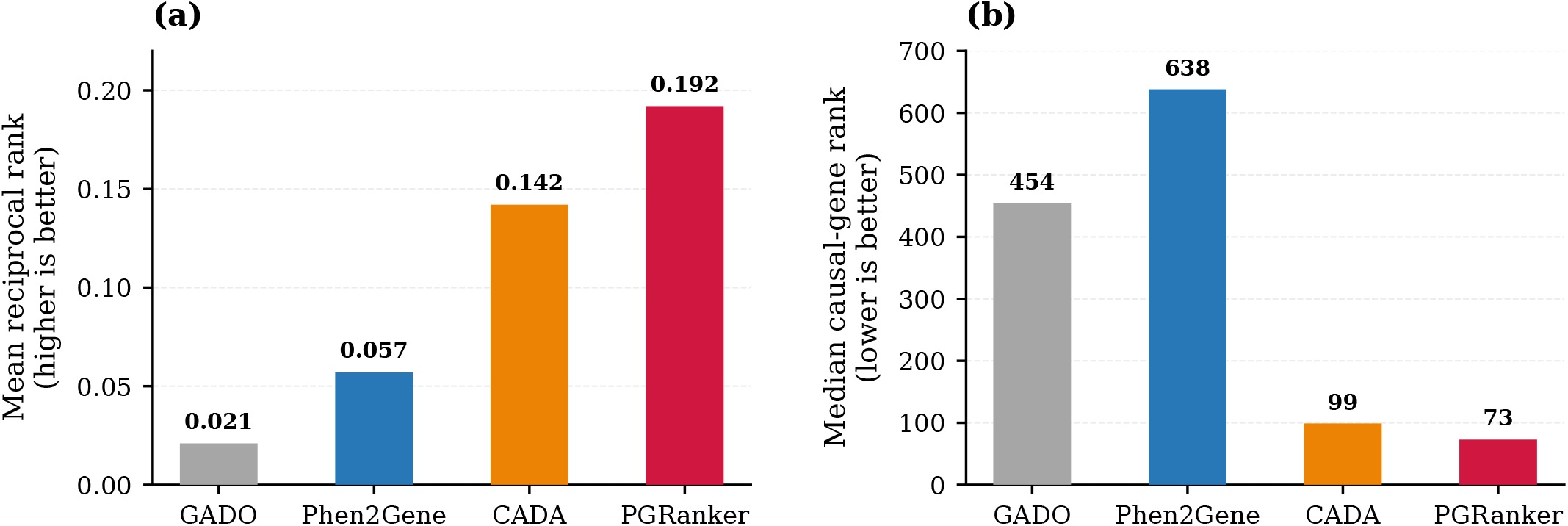
Mean reciprocal rank (MRR) and median rank of causal genes of all methods across the benchmark-277. Overall, PGRanker leads on both mean reciprocal rank (0.192) and median rank (73) of the causal gene averaged over 277 cases of the benchmark.

### Phenotype-gene evidence

Phenotype-gene evidence quantifies, for every candidate gene, the fraction of a case’s indexed HPO terms supported by curated annotation. Phenotype-gene associations obtained from HPO were encoded in the binary matrix *A* ∈ {0, 1}^|*ℋ*|*×N*^, where *A*_*hg*_ = 1 if HPO term *h* is associated with gene *g* and *A*_*hg*_ = 0 otherwise. The resulting matrix contains 557,803 associations.

For case *i* with 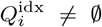, the phenotype-gene evidence support for candidate gene *g* is

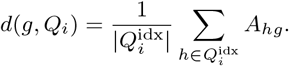

Terms outside ℋ contribute to neither the numerator nor the denominator. Evaluating this expression over all candidates yields the case-specific support vector

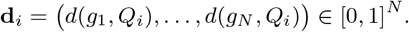

The lookup involves no learned parameters. Each gene’s scalar support is subsequently mapped through a shared learned projection to initialize its node representation in the gene graph encoder.

### Patient-specific gene graph encoder

PGRanker propagates phenotype-gene evidence over a fixed gene–gene interaction graph, GeneNet. The graph is shared across cases, but its node inputs are derived from each case’s phenotype profile, yielding patient-specific gene representations.

We constructed GeneNet from HumanNet-XC (v3) [12], whose *N* = 21,592 genes define the candidate universe *G*. The processed network contained 1,125,494 directed edge entries.

Reverse edges were added and duplicates merged by retaining the maximum confidence, yielding 2,250,988 directed edges. Confidences were min–max normalised to 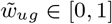. For each gene, we retained its 50 highest-confidence incoming edges and then restored symmetry, producing a final graph of 1,170,568 directed edges. Consequently, node degree can exceed 50.

For case *i*, each gene’s phenotype-gene evidence is mapped through a shared learned projection to a node representation of dimension *D* = 256:

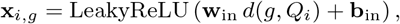

where **w**_in_, **b**_in_ ∈ ℝ^*D*^. All genes without direct support therefore start from the same vector.

A single graph-attention layer [28] propagates these inputs using *K* = 4 attention heads, whose outputs are averaged. Let *N* (*g*) denote the incoming neighbours of gene *g* in the final graph and 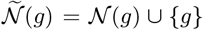 include its self-loop. For head *k*, the attention logit for a message from source gene 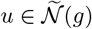 to target gene *g* is

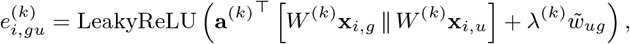

where *W* ^(*k*)^ ∈ ℝ^*D×D*^ is a learned transformation, **a**^(*k*)^ ∈ ℝ^2*D*^ is a learned attention vector, and ∥ denotes concatenation. The effective learned scalar coefficient *λ*^(*k*)^ controls the contribution of interaction confidence to the attention logit. Confidence thus modulates attention rather than scaling messages directly. Each self-loop receives the mean confidence of the gene’s incoming edges.

Attention coefficients are normalized over the target gene’s neighborhood:

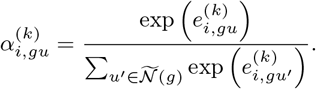

During training, dropout of 25% is applied to these normalized coefficients before aggregation. Let 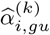 denote the coefficients after dropout during training and the unchanged coefficients 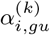 at inference. The weighted neighborhood messages are averaged across heads:

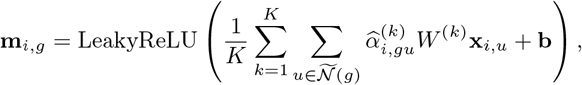

where **b** ∈ ℝ^*D*^ is a learned bias.

Finally, the graph-updated representation is obtained through a residual connection followed by layer normalization:

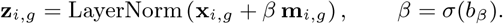

Here, *σ* is the sigmoid function, and *β* is a learned scalar residual weight shared across genes and cases, initialized to 0.10. This update retains a direct path from each gene’s own phenotype support while incorporating information from its interaction partners.

### Phenotype query encoder

The phenotype-gene evidence reduces a case’s indexed phenotype profile to one scalar per gene. The graph encoder propagates these support values but does not explicitly represent the identities of the contributing HPO terms. The phenotype query encoder represents the profile itself, operating on the same indexed terms 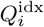 used for lookup.

Each term *h* ∈ ℋ is assigned a trainable embedding **e**_*h*_ ∈ ℝ^256^. An attention network with architecture 256 → 128 → 1 and a tanh hidden activation maps each embedding to a scalar

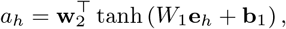

where *W*_1_ ∈ ℝ^128*×*256^ and **b**_1_, **w**_2_ ∈ ℝ^128^ are learned parameters. A shared scalar output bias is omitted because it cancels under softmax.

The logits are normalized over the terms present in the case:

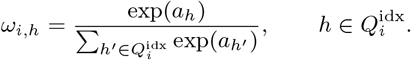

The phenotype-query representation is their attention-weighted embedding sum:

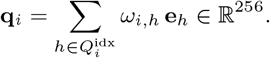

Each term’s logit depends only on its embedding, whereas its normalized weight depends on the other terms present through the softmax denominator. Pooling is invariant to term order and accommodates non-empty indexed profiles of varying size. The embeddings and attention parameters are learned end-to-end through the ranking objective, without separate supervision, and held fixed at inference. The encoder does not use the HPO ontology structure. The resulting query representation is shared across all candidate genes for a case and is combined with each gene’s graph-updated representation in the scorer.

### Integrated genome-wide scorer

For each patient *j* and candidate gene *g*_*i*_, PGRanker combines the graph-refined gene representation **z**_*ij*_ with the phenotype-query representation **q**_*j*_ to compute a single relevance score.

The two representations are concatenated together with their element-wise product and absolute difference,

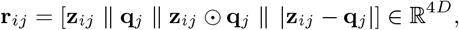

and a multilayer perceptron maps this joint representation to the scalar relevance score,

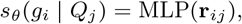

where *θ* denotes all learnable parameters of PGRanker.

The element-wise product and absolute difference capture agreement and mismatch between each gene’s representation and the patient’s phenotype profile. Because **q**_*j*_ is identical for all candidate genes of patient *j*, it does not reorder genes by itself but influences the ranking through its interaction with **z**_*ij*_. The scorer is applied to every candidate gene *g*_*i*_ ∈ G, and genes are ranked in descending order of *s*_*θ*_(*g*_*i*_ | *Q*_*j*_).

### Model training

PGRanker was trained on Mendelian disorders from OMIM [1]using disease phenotype profiles from HPO [24], restricted to OMIM records and phenotypic abnormalities, with negated annotations excluded. Disorders were split by unique OMIM identifier in a 70:15:15 ratio, keeping all gene associations of a disorder in one partition. Restricting each partition to disorders with at least one term in ℋ and one causal gene in *G* yielded 4,208 training, 904 validation and 887 test queries.

Each training occurrence used a disorder’s full indexed HPO set 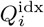, one selected causal gene *g*^+^, and a set *C*_*i*_ of 1,024 distinct negative genes, excluding all genes associated with that disorder. Negatives were sampled from genes with non-zero direct support, interaction partners of *g*^+^ and the full candidate universe, providing difficult alternatives alongside randomly sampled candidates.

To reduce reliance on direct support at causal genes, the support values of all known causal genes for the disorder were set to zero for 20% of training queries. This masking affected both the graph input and the auxiliary direct-support branch, while leaving the HPO query unchanged. Half of each epoch was sampled from queries containing a causal gene with no direct support but at least one supported interaction partner. We determined eligibility for this sampling procedure using the unpruned network, whereas graph propagation used the pruned GeneNet graph. Both masking and enriched sampling were applied during training only.

Training used a combined score comprising an auxiliary direct-support score *l*_*i*_(*g*) and the graph-based scorer output *r*_*i*_(*g*), weighted by a learned adaptive gate *γ*_*i*_(*g*):

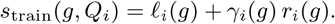

Here, *r*_*i*_(*g*) denotes the score from the genome-wide scorer described above. Reported inference used *r*_*i*_(*g*) alone.

For each query, a softmax cross-entropy ranking loss was evaluated over the positive gene and the 1,024 sampled negatives:

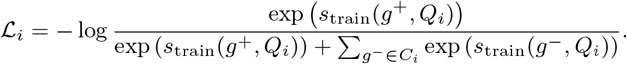

Gradients from this objective jointly updated the support projection, graph-attention encoder, HPO term embeddings, query-attention network, genome-wide scorer, and auxiliary branch and gate. The association matrix *A* and gene network remained fixed; case-specific support was computed from *A* and modified only by the training-time masking described above.

Because node inputs differ between cases, we performed a full-graph forward pass for each query and accumulated gradients over 32 queries before each optimizer step. After every epoch, the model ranked the complete gene universe for each validation disorder using *s*_train_. The checkpoint with the highest recall@50 was retained, with MRR as a tie-breaker. Reported metrics were averaged across three models trained with independent random seeds. Supplementary Section 2 provides further training and model configuration details.

### Benchmarks and baseline methods

We compared PGRanker with three phenotype-driven gene-prioritization methods that accept HPO terms without variant information: Phen2Gene, which aggregates curated phenotype– gene associations; CADA, which performs prioritization using embeddings of an HPO–gene knowledge graph; and GADO, which predicts phenotype–gene associations from gene co-regulation. Each method received the case’s observed HPO terms and returned a ranked candidate-gene list. Baseline versions, resource bundles and execution settings are provided in Supplementary Section 3.

We evaluated all four methods on two cohorts of solved clinical cases, each with an observed HPO profile and one confirmed causal gene. Benchmark-277 is the multi-cohort evaluation set distributed with Phen2Gene [29]. It originally contains 281 cases from five independently ascertained cohorts; excluding four cases without HPO terms left 277 cases, with a median of nine terms per case (range 1–56). The second cohort comprises 21 solved rare-disease cases spanning 20 distinct causal genes, with sparse profiles of one to four HPO terms per case. Three underlying gene–disease relationships were previously published [23, 22, 26]; the remaining 18 cases are unpublished. Further details of the benchmark datasets, baseline methods, and PGRanker implementation details are provided in Supplementary Sections 1-3.

#### Evaluation metrics and statistical analysis

For each of the *M* evaluated cases, we recorded the rank *r*_*i*_ of its confirmed causal gene. Cases for which a method returned no usable ranking, or did not return the causal gene, were treated as misses and retained in the benchmark denominator. Recall@*k* is the fraction of cases whose causal gene was ranked within the top *k*, and mean reciprocal rank (MRR) the mean of 1*/r*_*i*_ over all cases, with misses contributing zero to both. As a measure less influenced by extreme ranks we additionally report the median causal-gene rank, computed under a fixed denominator in which an unranked causal gene receives the sentinel rank *N* + 1 = 21,593, one position below the complete candidate universe.

PGRanker comprises three models trained with three different seeds, each of which ranks all candidate genes independently; rank, reciprocal rank, and recall@*k* were computed per seed and averaged per case, and all reported statistics were derived from these seed-averaged case values. Reported PGRanker ranks are therefore means over three runs rather than the output of a single model, and the recall@*k* reported is the mean over seeds of the fraction of cases recovered within the top *k*. Baseline rankings were evaluated in their returned order after gene-symbol normalisation and removal of duplicate entries.

We estimated 95% confidence intervals for recall@*k* and MRR by percentile bootstrap resampling at the case level, drawing 2,000 resamples of *M* cases with replacement and taking the 2.5th and 97.5th percentiles of the resulting distribution; the same resampled case indices were used across methods, preserving the paired benchmark design. Pairwise differences between PGRanker and each baseline were assessed on the per-case reciprocal ranks with a one-sided Wilcoxon signed-rank test under the alternative that PGRanker produced higher reciprocal ranks, retaining zero paired differences by the Pratt procedure and adjusting the three resulting *p*-values by the Holm procedure. Cohort-specific Recall@10 values were calculated descriptively within each cohort using the same fixed-denominator definition, without separate cohort-level tests or confidence intervals.

## Results

### PGRanker outperforms existing Mendelian ranking methods

PGRanker achieved the highest mean reciprocal rank and lowest median causal-gene rank on Benchmark-277 (Table 1). Its MRR was 0.192, compared with 0.142 for CADA, 0.057 for Phen2Gene and 0.021 for GADO. The median causal-gene rank was 73, compared with 99 for CADA, 638 for Phen2Gene and 454 for GADO.

**Table 1.** Ranking performance on Benchmark-277. We report the recall@k, mean reciprocal rank (MRR), and the median causal gene rank across the 277-case benchmark for all four methods. P-values were computed using a one-sided Wilcoxon signed-rank test on per-case reciprocal ranks versus PGRanker (PGR) and Holm-corrected across the three baselines. We observe that PGRanker achieves the best overall recall@*k* across all values of *k*, particularly recovering the causal gene in its top-1 ranked list in 14% of cases. PGRanker also achieves the lowest median causal gene rank, most consistently ranking the causal gene highly.

| Metric | GADO | Phen2Gene | CADA | PGRanker (PGR) |
| --- | --- | --- | --- | --- |
| MRR | 0.021 | 0.057 | 0.142 | **0.192** |
| Recall@1 | 0.40% | 2.20% | 9.00% | **14.00%** |
| Recall@5 | 2.20% | 9.00% | 19.90% | **24.20%** |
| Recall@10 | 5.80% | 11.90% | 23.80% | **27.90%** |
| Recall@20 | 7.90% | 18.80% | 30.00% | **33.20%** |
| Recall@50 | 15.20% | 26.00% | 40.40% | **44.80%** |
| Recall@100 | 21.30% | 31.40% | 50.20% | **54.00%** |
| Median rank | 454 | 638 | 99 | **73** |
| Wilcoxon vs. PGR ($p_{\text{Holm}}$) | $4.35 \times 10^{-27}$ | $8.74 \times 10^{-15}$ | $5.10 \times 10^{-3}$ | — |

Recall@ $k$ denotes the percentage of cases for which the causal gene was ranked within the top $k$ candidates. Lower median rank (of the causal gene) indicates better performance. PGR metrics are reported as the average over 3 seeds, with the median rank rounded to the nearest integer. Boldface indicates the best-performing method for each metric.

PGRanker also achieved the highest Recall@*k* at every evaluated cutoff from 1 to 100 (Figure 3). Compared with CADA, the strongest baseline, Recall@1 increased from 9.0% to 14.0%, an improvement of 5.0 percentage points. Recall@5 increased from 19.9% to 24.2% and Recall@10 from 23.8% to 27.9%. The advantage persisted at broader cutoffs, with Recall@50 of 44.8% versus 40.4% and Recall@100 of 54.0% versus 50.2%.

**Figure 3.**
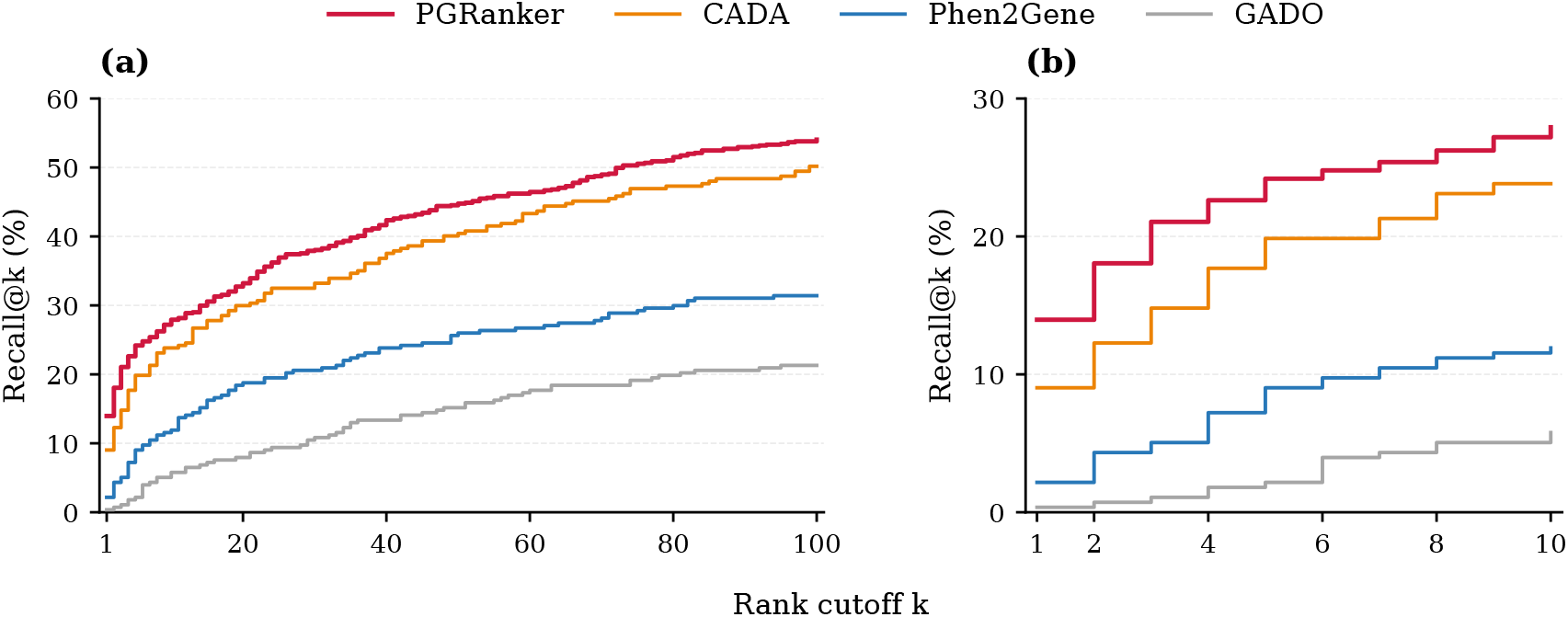
Recall@*k* of all four methods on benchmark-277 representing the percentage of cases each method was able to rank the causal gene of the case in its top *k* ranked list of genes. We observe that PGRanker leads in recovering the causal gene at every cutoff of K, most notably in its top-1 rank for 14% of cases compared to 9% for the best baseline.

Per-case reciprocal-rank comparisons favored PGRanker over all three baselines after Holm correction, including CADA (*p*_Holm_ = 5.1 × 10^*™*3^; Table 1). Its higher MRR and recall therefore occurred despite returning causal genes for fewer cases than Phen2Gene and GADO, demonstrating stronger prioritization near the top of the ranking rather than greater overall coverage.

### Performance under phenotype degradation

Clinical phenotype profiles may be incomplete, contain observations unrelated to the causal disorder, or use terms of differing specificity. We therefore evaluated four forms of input degradation: removal of observed terms, addition of unrelated incorrect terms, substitution with incorrect terms, and replacement by less-specific ancestors. The analysis used the 231 Benchmark-277 cases containing at least five HPO terms, with model parameters and annotation resources held fixed. Random perturbations were repeated 20 times per case and level, whereas ancestor replacement was deterministic. Unperturbed Recall@10 on this subset was 31.2% for PGRanker, compared with 26.0% for CADA, 12.6% for Phen2Gene and 6.9% for GADO.

PGRanker achieved the highest Recall@10 at every tested removal, addition and substitution level (Figure 4a–c). Its Recall@10 AUC was 25.86% under removal, 27.63% under addition and 19.30% under substitution, compared with 21.29%, 24.15% and 15.82%, respectively, for CADA, the strongest baseline in these regimes. These correspond to advantages of 4.57, 3.48 and 3.48 percentage points. Substitution caused a pronounced decline, with all methods recovering few causal genes within the top ten at the highest tested severity.

**Figure 4.**
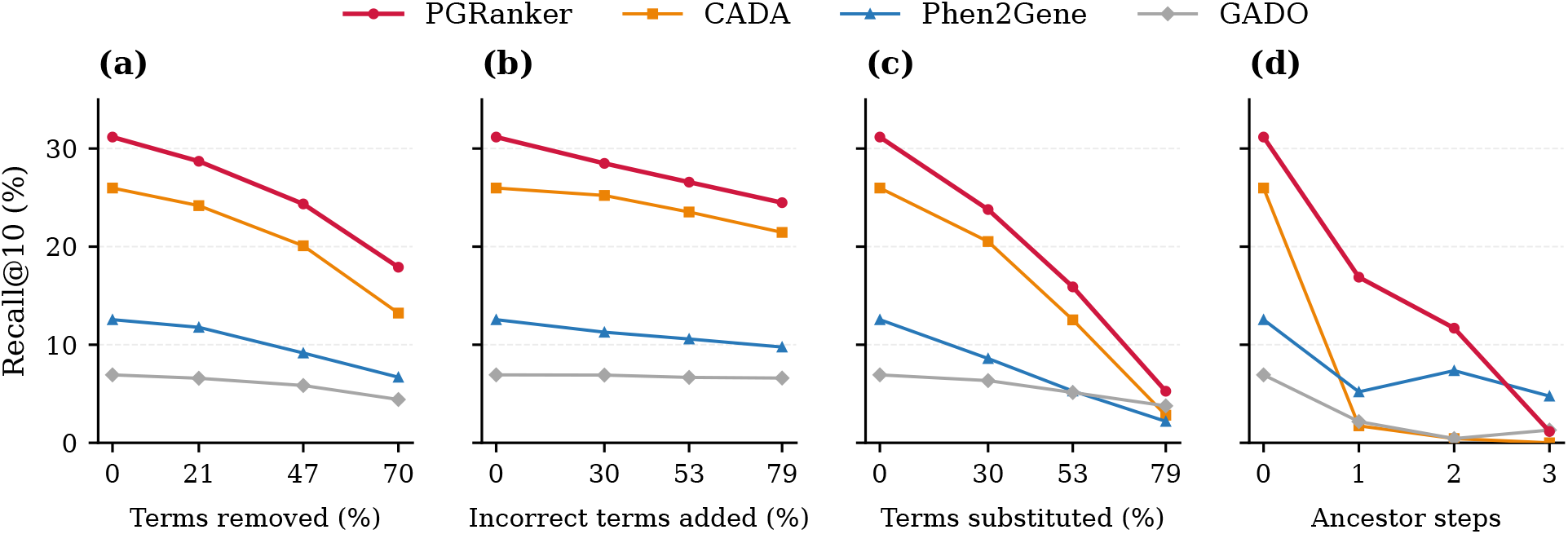
Robustness to phenotype degradation on Benchmark-277. Recall@10 across increasing severity of four phenotype-degradation settings in the 231 cases containing at least five HPO terms: (**a**) percentage of observed terms removed, (**b**) percentage of unrelated incorrect terms added, (**c**) percentage of observed terms substituted with incorrect terms, and (**d**) number of ancestor-replacement steps. Random perturbations were averaged over 20 repetitions per case and level, whereas ancestor replacement was deterministic.

**Figure 5.**
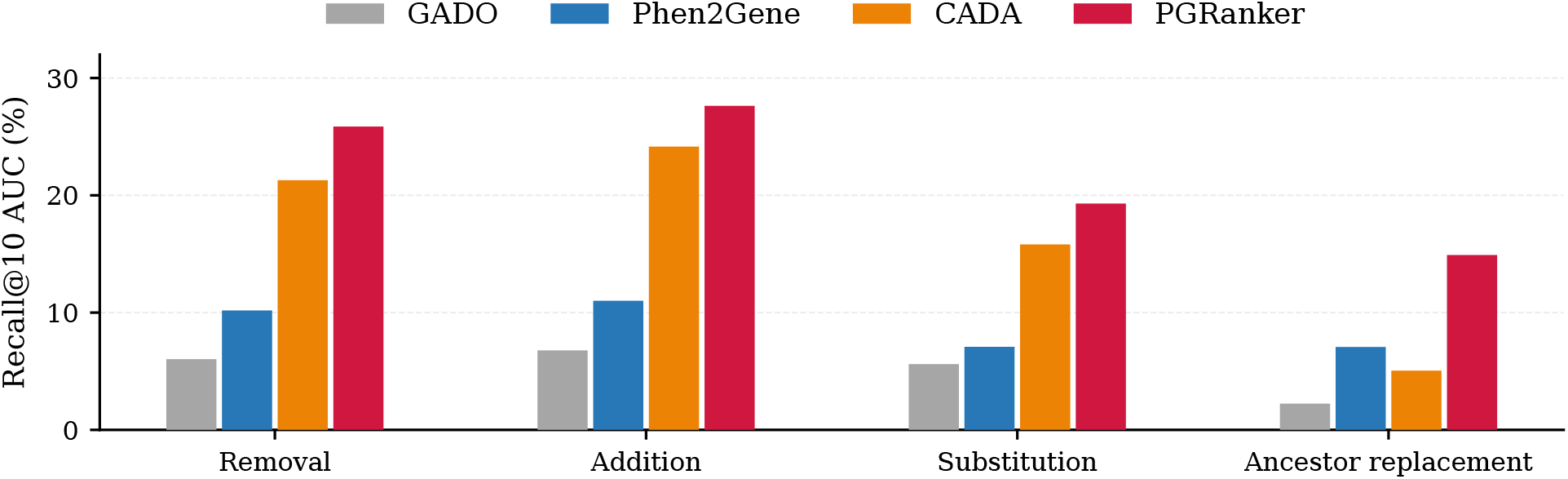
Overall robustness across phenotype-degradation settings. Area under the Recall@10 trajectory (AUC) over normalized perturbation severity for term removal, addition of unrelated incorrect terms, substitution with incorrect terms, and replacement by less-specific ancestor terms. Higher AUC indicates better overall Recall@10 across the tested degradation range. For ancestor replacement, CADA and GADO rejected 293 and 273 of the 693 perturbed queries, respectively, because the ancestor identifiers are absent from their bundled ontologies; these queries were retained as misses.

Ancestor replacement produced the largest AUC advantage over the next-best method: 14.91% for PGRanker versus 7.07% for Phen2Gene, a difference of 7.84 percentage points. PGRanker led after one and two replacement steps, whereas Phen2Gene led after three (Figure 4d). Comparisons with CADA and GADO additionally reflect ontology-version compatibility: these methods rejected 293 and 273 of the 693 perturbed queries, respectively, because ancestor identifiers were absent from their bundled ontologies. These queries were retained as misses.

PGRanker also achieved the highest MRR AUC in all four regimes. Together, these results show higher absolute ranking performance across the tested degradation trajectories, while severe phenotype substitution and loss of specificity substantially reduced performance (additional results are provided in Supplementary Section 4.1).

### Contribution of phenotype-gene evidence and learned refinement

We compared the full model with phenotype-gene evidence and a configuration without seeding on Benchmark-277 (Table 2). Phenotype-gene evidence alone was used to rank genes by their support scores *d*(*g, Q*_*i*_), while the unseeded configuration omitted this support from the graph input.

**Table 2.** PGRanker architecture ablation. Integrating the GeneNet component via seeding from direct lookup improves overall ranking performance on benchmark-277, though it catastrophically fails on its own. The direct lookup mechanism is by far the most essential component of the model. All results are reported as the average over three independently seeded models; the median rank is rounded to the nearest integer.

| Configuration | MRR | R@1 | R@5 | R@10 | R@20 | R@50 | R@100 | Median rank |
| --- | --- | --- | --- | --- | --- | --- | --- | --- |
| GeneNet (without seed) | 0.0006 | 0.00% | 0.00% | 0.12% | 0.12% | 0.24% | 0.60% | 6722 |
| Phenotype-gene evidence only | 0.1786 | 12.64% | 22.74% | **28.16%** | **33.94%** | 42.24% | 52.71% | 87 |
| Full model | **0.1918** | **13.96%** | **24.19%** | 27.92% | 33.21% | **44.77%** | **54.03%** | **73** |

Phenotype-gene evidence supplied the dominant ranking signal. Direct lookup alone achieved an MRR of 0.179 and a median causal-gene rank of 87, whereas the configuration without lookup seeding achieved an MRR below 0.001 and a median rank of 6,722. Building on this strong lookup baseline, the full model increased MRR to 0.192 and improved the median rank to 73. Recall@1 increased from 12.6% to 14.0%, Recall@5 from 22.7% to 24.2%, Recall@50 from 42.2% to 44.8%, and Recall@100 from 52.7% to 54.0%. Direct lookup retained slightly higher Recall@10 and Recall@20, at 28.2% and 33.9%, compared with 27.9% and 33.2% for the full model.

Case-level comparisons illustrate improvements to both strong lookup predictions and candidates initially outside the top 50 (Table 3). The full model promoted *DHX30* from rank 4 to 1 and *TP53* from 3 to 1, while *TBC1D23* and *AHDC1* both moved from rank 10 to 2. It also brought *RAC1* from rank 62 to 4 and *SAMHD1* from 261 to 42, placing both within the top 50 and *RAC1* within the top five. Similarly, *SYNGAP1* improved from 72 to 18, *MRPS34* from 40 to 22, *KAT6B* from 62 to 26, and a second *DHX30* case from 98 to 28. These changes substantially reduced the number of candidates preceding the causal gene.

**Table 3.** Causal gene rank improvement with GAT. The seeded GeneNet can enhance the ranking performance of direct lookup, in some cases drastically by rescuing missed genes ranked outside of the clinically relevant range within the top 50 or higher.

| Causal gene | Lookup rank | PGR rank | Change |
| --- | --- | --- | --- |
| DHX30 | 4 | **1 (+3)** | +75.0% |
| TP53 | 3 | **1 (+2)** | +66.7% |
| TBC1D23 | 10 | **2 (+8)** | +80.0% |
| AHDC1 | 10 | **2 (+8)** | +80.0% |
| RAC1 | 62 | **4 (+58)** | +93.5% |
| SYNGAP1 | 72 | **18 (+54)** | +75.0% |
| MRPS34 | 40 | **22 (+18)** | +45.0% |
| KAT6B | 62 | **26 (+36)** | +58.1% |
| DHX30 | 98 | **28 (+70)** | +71.4% |
| SAMHD1 | 261 | **42 (+219)** | +83.9% |

The benefits varied across cases (Table 4). For *ACTG2* and *RAB11B*, the full model moved the causal gene down by one position, from 3 to 4 and from 9 to 10, respectively. It also worsened the already low rankings of *CAPN5* and *MYH10*. Conversely, *NAA15* improved by 2,526 positions, from 14,608 to 12,082, but remained outside the top 100. These examples distinguish numerical rank improvements from recovery into a short candidate list.

**Table 4.** GeneNet sometimes fails to rescue or even worsens ranking performance of direct lookup. GeneNet sometimes worsens good rankings from direct lookup, often consistently across cases for the same causal gene. GeneNet is highly dependent on the direct lookup seeding and thus even in some cases where it improves catastrophic failures cannot reliably rescue these genes by re-ranking them within the top-50.

| Causal gene | Lookup rank | PGR rank | Change |
| --- | --- | --- | --- |
| ACTG2 | 3 | 4 (-1) | -33.3% |
| RAB11B | 9 | 10 (-1) | -11.1% |
| CAPN5 | 1274 | 2127 (-853) | -67.0% |
| MYH10 | 5716 | 6654 (-938) | -16.4% |
| NAA15 | 14608 | 12082 (+2526) | +17.3% |

Together, these results demonstrate the value of using phenotype-gene evidence to initialize the learned ranking model. The full model improves MRR, median rank, and four of the six reported recall cutoffs over direct lookup, with individual cases showing both refinement into the first few positions and recovery into the top 50.

### Mixed rare diseases cohort case study

To assess generalisation to genuinely unseen patients, we evaluated all four methods on a clinical cohort of 21 mixed monogenic rare-disease cases, most of them unpublished. These cases span phenotypically unrelated disorders and are annotated sparsely. As for Benchmark-277, each method received only the observed HPO terms and was scored against the confirmed causal gene. Further details of the mixed rare diseases benchmark are provided in Supplementary Section 1.3. All methods performed substantially worse than on Benchmark-277. PGRanker attained the highest recall at every cutoff and was the only method with non-zero Recall@1 and Recall@5. Its Recall@100 was 50.8%, compared with 33.3% for CADA, the strongest baseline (Table 5).

**Table 5.** Performance comparison on the 21-case causal-gene benchmark. Recall@*k* denotes the proportion of cases for which the causal gene was ranked within the top *k* candidates. Lower median causal-gene rank and higher average recall of the causal gene indicate better performance.

| Method | Median Rank | Recall@1 | Recall@5 | Recall@10 | Recall@20 | Recall@50 | Recall@100 |
| --- | --- | --- | --- | --- | --- | --- | --- |
| PGRanker | **123** | **4.80%** | **6.30%** | **11.10%** | **22.20%** | **38.10%** | **50.80%** |
| Phen2Gene | 252 | 0.00% | 0.00% | 0.00% | 4.80% | 9.50% | 19.00% |
| CADA | 134 | 0.00% | 0.00% | 4.80% | 4.80% | 28.60% | 33.30% |
| GADO | 423 | 0.00% | 0.00% | 0.00% | 0.00% | 9.50% | 9.50% |

PGRanker also achieved the best median causal rank of 123 compared to the best baseline CADA at 134, and substantially led against Phen2Gene and GADO at 252 and 423, respectively. Detailed results for each individual case are provided in Supplementary Section 4.2.

## Discussion

PGRanker integrates phenotype–gene evidence with patient-specific gene-network representations and an attention-pooled phenotype query to rank 21,592 candidate genes from HPO terms alone. Its central design choice is to use each patient’s phenotype support to initialise the gene graph before scoring, allowing the learned representations to combine local support with evidence from interaction partners. On Benchmark-277, PGRanker achieved the highest MRR, lowest median causal-gene rank, and highest Recall@*k* at every evaluated cutoff from 1 to 100. The increase in Recall@1 from 9.0% for CADA to 14.0% for PGRanker demonstrates an improvement at the first-ranked candidate, while gains through Recall@100 show that the advantage extends beyond the highest-priority positions.

The comparison with direct lookup helps explain the contribution of this integration. Phenotype-gene evidence alone provided a strong baseline, with an MRR of 0.179 and a median causal-gene rank of 87. The full model improved these to 0.192 and 73, respectively, and increased recall at four of the six tabulated cutoffs. Thus, learned refinement adds value even when explicit annotation support already provides an informative ranking. The poor performance of the configuration without lookup seeding further emphasises the importance of supplying patient-specific evidence to the graph. Because the full model also includes phenotype-query encoding and learned scoring, its improvement over direct lookup reflects the combined refinement rather than an isolated measurement of message passing.

The degradation experiments extend this finding to incomplete and corrupted phenotype profiles. PGRanker achieved the highest Recall@10 and MRR AUC in all four regimes and retained the highest Recall@10 at every tested removal, addition and substitution level. Under ancestor replacement, it led after one and two steps, whereas Phen2Gene led after three. These results demonstrate sustained absolute ranking performance across a range of input quality, with severe substitution and loss of specificity remaining challenging. Comparisons involving CADA and GADO under ancestor replacement also reflect incompatibility between generated terms and their bundled ontology versions, highlighting the importance of distinguishing phenotype degradation from resource compatibility.

The 21-case in-house cohort provides a complementary evaluation on diverse monogenic disorders described by only one to four HPO terms per case. The current aggregate results favor PGRanker at every reported recall cutoff and in median causal-gene rank, supporting its applicability to sparse clinical profiles. Larger cohorts with documented ascertainment and phenotype curation would allow a more precise assessment of performance across profile sizes and disease groups.

Several aspects of the evaluation suggest useful next steps. The methods use different annotation resources and releases, so their comparison reflects both modelling choices and available knowledge. Publication dates alone do not establish equivalent annotation coverage, and unpublished patients may have disease–gene associations already represented in these resources. Evaluation using harmonised annotation snapshots and temporally separated clinical cohorts would help distinguish these effects. Training on complete OMIM disease profiles also differs from application to individual patient profiles; incorporating realistic phenotype incompleteness and noise during training is a natural extension of the present robustness experiments.

At the model level, direct support weights indexed phenotype terms equally, and a single graph-attention layer limits propagation to immediate neighbors within the HumanNet-derived candidate universe. Future work could examine specificity-aware support, alternative interaction networks, and selective multi-hop propagation. Component-controlled ablations would help identify which learned operations contribute most to the observed gains, while confidence-aware refinement could preserve strong lookup predictions and target cases that benefit from re-ranking. Variant-level evidence offers a further route to integrating phenotype-driven prioritization into diagnostic workflows.

Overall, PGRanker demonstrates that phenotype-gene evidence can serve as an effective foundation for patient-specific network learning. Combining that evidence with learned gene and phenotype representations improves genome-wide ranking and maintains strong performance across varied phenotype-input conditions, providing a practical basis for prioritizing causal genes from clinical phenotype descriptions.

## Data and code availability

All datasets and code for PGRanker are publicly available at: https://github.com/ivelet/PGRanker.

## Author contributions statement

R.K. designed, developed, and implemented the model and pipeline, designed and conducted all experiments and analyses, and wrote the manuscript. M.S. provided clinical data and manual exome data analyses, contributed to discussions, and revised the manuscript. A.F. and A.K. contributed to discussions and revised the manuscript. V.D.T. supervised the study, contributed to discussions and revised the manuscript. R.B. contributed to discussions, revised the manuscript, supervised the study, and provided funding. All authors contributed to discussions and reviewed the final manuscript.

## Funding and Acknowledgments

This research was funded by the Deutsche Forschungsgemeinschaft (DFG, German Research Foundation) under grant number 539134284, through EFRE (FEIH 2698644) and the state of Baden-Württemberg; Deutsche Forschungsgemeinschaft (DFG, German Research Foundation) [Project-ID 499552394 – SFB 1597]; and the state of Baden-Württemberg through bwHPC and the German Research Foundation (DFG) [grant number INST 35/1597-1 FUGG].

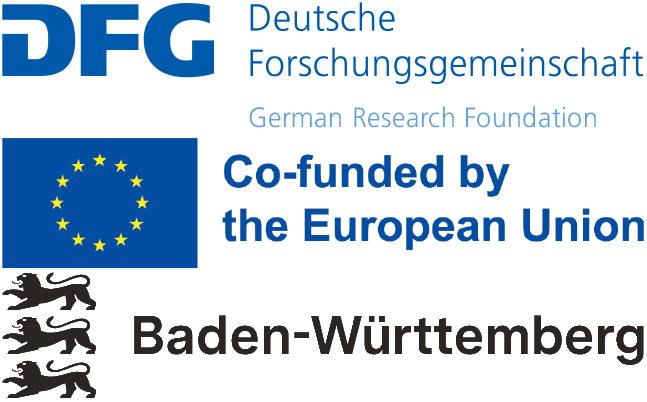

