## Supplement for "PGRanker: Enhancing Phenotype Knowledge with Graph Learning for Mendelian Gene Prioritization"

PGRanker: Patient-Specific Phenotype-Guided Graph Learning for Mendelian Gene Prioritization

#### S1 Datasets and preprocessing

##### S1.1 PGRanker source datasets and preprocessing

PGRanker uses phenotype and disease–gene information from the Human Phenotype Ontology (HPO) and OMIM, together with gene–gene interactions from HumanNet-XC (v3). Phenotype-gene evidence was obtained from HPO ('phenotype_to_genes.txt' release 2026-06-23). The candidate-gene universe $\mathcal{G}$ comprises the $N = 21{,}592$ genes in the processed HumanNet-XC network. The indexed phenotype vocabulary $\mathcal{H}$ contains the 8,419 HPO terms annotated to OMIM disorders across the training, validation and test partitions.

Disease phenotype profiles were obtained from HPO, restricted to OMIM records and phenotypic abnormalities, with negated annotations excluded. HPO identifiers were standardized by removing surrounding whitespace, converting them to uppercase and replacing underscores with colons. Valid identifiers were deduplicated and restricted to the indexed phenotype vocabulary. Alternative and obsolete identifiers were not explicitly remapped during PGRanker's standard profile preprocessing.

HPO–gene associations were obtained from `phenotype_to_genes.txt` and represented in the binary matrix $A \in \{0,1\}^{|\mathcal{H}|\times N}$ over indexed terms and candidate genes. The matrix contains 557,803 distinct term–gene associations; repeated entries for the same pair do not increase its binary value.

The processed HumanNet-XC network contained 1,125,494 directed edge entries. Reverse edges were added and duplicates merged by retaining the maximum confidence, yielding 2,250,988 directed edges. Confidence scores were min–max normalised to $\tilde{w}_{uv} \in [0, 1]$. For each gene, the 50 highest-confidence incoming edges were retained, after which symmetry was restored, resulting in 1,170,568 directed edges. Consequently, the final node degree can exceed 50.

##### S1.2 Benchmark-277

We used the real-patient evaluation set distributed with Phen2Gene, comprising 281 solved monogenic cases, each with an HPO phenotype profile and one confirmed causal gene [1]. Four cases from the AJHG cohort lacked HPO terms and were excluded, leaving the 277 cases evaluated here.

The benchmark combines five independently ascertained cohorts that differ in provenance and phenotype curation:

- **TAF1** comprises 14 cases from a series describing *TAF1*-associated dysmorphism, intellectual disability and neurological manifestations, with physician-curated HPO terms [2].
- **ColumbiaU** comprises 27 cases with 24 distinct causal genes from a deep-phenotyping study at Columbia University Medical Center. HPO terms were curated manually from clinician-defined phenotypes in electronic health records [3].

- **DGD** comprises 85 cases with 75 distinct causal genes from the Department of Genomic Diagnostics at the Children's Hospital of Philadelphia. These de-identified diagnostic cases have physician-curated HPO terms [4].
- **CSH** comprises 72 cases with 59 distinct causal genes assembled from 61 *Cold Spring Harbor Molecular Case Studies* reports.
- **AJHG** comprises 79 retained cases from an original collection of 83 cases, assembled from 14 *American Journal of Human Genetics* reports.

For CSH and AJHG, HPO terms were extracted from article text using Doc2HPO [5] and manually reviewed to remove negated and duplicated terms [1]. Several patients within these literature cohorts share a newly implicated causal gene.

Each case has one confirmed causal gene. All evaluated methods received only the observed HPO terms, without variant-level evidence or a pre-filtered candidate-gene list. Across Benchmark-277, the number of input HPO terms ranged from 1 to 56, with a median of nine and an interquartile range of 5–14. These counts describe the supplied profiles, before method-specific vocabulary filtering.

##### S1.3 Mixed rare-disease cohort

This cohort comprises 21 solved cases spanning 20 distinct causal genes across phenotypically diverse monogenic disorders. Cases were drawn from a mix of rare-disease genetic diagnostics and gene-discovery programmes comprised of both published and unpublished work. Results from manual exome data analyses for cases 1,9, 10,11 and12 have been previously published (PMID:33764426, PMID:30760291, PMID:32282878, PMID:30520571, PMID:30487245).

Three underlying gene–disease relationships were previously reported in gene-discovery studies:

- *PPP1R21* was implicated in a neurodevelopmental syndrome involving hypotonia, developmental delay, facial dysmorphism and thinning of the corpus callosum. The published study identified four homozygous truncating alleles in four independent families, localised the protein to the early endosome and demonstrated delayed transferrin clearance in patient fibroblasts [6].
- *MAB21L1* was implicated in an autosomal recessive syndrome involving a characteristic facial gestalt, corneal dystrophy, cerebellar and genital anomalies (COFG syndrome). The published study identified four loss-of-function variants and one missense variant in ten affected individuals from five unrelated consanguineous families [7].
- *VPS41*, encoding a subunit of the HOPS tethering complex, was implicated in a progressive neurodevelopmental disorder involving cognitive impairment, cerebellar atrophy, ataxia, dystonia and nystagmus. The published study identified homozygous variants in nine affected individuals from five unrelated families, with zebrafish modelling supporting a lysosomal mechanism [8].

The remaining benchmark cases are unpublished solved cases from the programme. For all cases, monogenic causes of well established human disease genes were manually identified fitting the cases phenotypes.

Each case has one confirmed causal gene. Methods received the observed HPO terms without variant information or a pre-filtered candidate-gene list, allowing evaluation across diverse disorders with sparse phenotype descriptions. Ethics approval was granted by The Freiburg University Ethics committee (approval 22-1413-S1).

### S2 Implementation details

#### S2.1 Model configuration

The projected gene inputs, graph-updated gene representations and phenotype-query representations have dimension $D = 256$. The gene graph encoder uses one graph-attention layer with four heads whose outputs are averaged. The support projection and graph aggregation use LeakyReLU activations with negative slope 0.01. Attention-coefficient dropout is 0.25 during training, and the learned scalar residual weight $\beta$ is initialised to 0.10.

The phenotype query encoder uses 256-dimensional trainable HPO-term embeddings and a $256 \to 128 \to 1$ attention network with a tanh hidden activation and a softmax over the terms present in the query. The genome-wide scorer maps the concatenated gene–query features through an MLP with dimensions $1{,}024 \to 512 \to 256 \to 1$. Both hidden layers use LeakyReLU activations with negative slope 0.01, followed by dropout with probabilities 0.25 and 0.125, respectively; the final linear layer produces a scalar ranking score. During training, an auxiliary direct-support branch projects each gene's scalar phenotype support to 256 dimensions and concatenates it with the phenotype-query representation before scoring it through a 512→512→256→1 MLP. A candidate-specific gate, implemented as a 513→128→1 network with LeakyReLU and a sigmoid output, receives the graph-updated gene representation, phenotype-query representation and scalar direct support. The combined training score adds the direct-support score to the gated graph score. The gate is initialized to 0.05. With probability 0.20 per training query, support for all known positive genes is masked to zero across the support-dependent pathways. At inference, PGRanker uses the ungated graph score.

#### S2.2 Optimisation and execution

A full-graph forward pass was performed for each query, and gradients were accumulated over 32 queries before each optimiser step. Optimisation used AdamW with a learning rate of $10^{-3}$, weight decay of $10^{-4}$ and gradient clipping at a maximum norm of 5, with BF16 mixed precision. Training ran for at most 100 epochs, with early stopping after 15 epochs without improvement in validation Recall@50. Checkpoint selection and the distinction between training and inference scores are described in the main-text Model training subsection. Three models were trained with independent random seeds: 42, 43, and 44. Each query was paired with one positive gene and 1,024 distinct negatives, with requested allocations of 256 supported genes, 256 HumanNet neighbors of the selected positive, 256 additional supported genes and 256 uniformly sampled genes. Sampling excluded all known positive genes and previously selected negatives. Pool shortfalls were filled by uniform sampling from the remaining candidate universe. PGRanker was implemented using PyTorch 2.3.1 and PyTorch Geometric 2.5.3 and trained on a single NVIDIA H200.

### S3 Baseline methods

We compared PGRanker with three phenotype-driven gene-prioritisation methods that accept HPO terms without variant information: Phen2Gene [1], CADA [9] and GADO [10]. These methods use curated phenotype associations, knowledge-graph embeddings and gene co-regulation, respectively.

#### S3.1 Phen2Gene

Phen2Gene aggregates weighted HPO-to-gene associations from the HPO-to-gene knowledge base (H2GKB), combining evidence obtained for individual phenotype terms. It was run at commit

Table S1: PGRanker model architecture and training configuration.

| Category | Setting | Value Actually Used |
|---|---|---|
| Model | Candidate gene pool size | 21,592 |
| Model | Indexed HPO terms | 8,419 |
| Model | Common representation dimension | 256 |
| Gene graph | Final directed edges | 1,170,568 |
| Gene graph | Edge selection | Top-50 incoming edges, restored symmetry |
| Gene graph | Edge confidence | Min–max normalized; used in attention |
| Gene graph | Support projection | Linear $1 \rightarrow 256$, LeakyReLU |
| Gene graph | GAT layers | 1 |
| Gene graph | GAT heads | 4, averaged |
| Gene graph | Self-loops | Enabled |
| Gene graph | Post-aggregation activation | LeakyReLU (slope = 0.01) |
| Gene graph | Attention-logit activation | LeakyReLU (slope = 0.2) |
| Gene graph | Attention dropout | 0.25 |
| Gene graph | Residual weight | Trainable; initialized to 0.10 |
| Gene graph | Normalization | LayerNorm |
| HPO encoder | HPO embedding dimension | 256 |
| HPO encoder | Attention network | $256 \rightarrow 128 \rightarrow 1$ |
| HPO encoder | Attention activation | Tanh, term-wise softmax |
| Scorer | Gene–query feature dimension | 1,024 |
| Scorer | MLP dimensions | $1{,}024 \rightarrow 512 \rightarrow 256 \rightarrow 1$ |
| Scorer | Hidden activation | LeakyReLU (slope = 0.01) |
| Scorer | Hidden-layer dropout | 0.25, 0.125 |
| Scorer | Inference score | Ungated graph-branch score |
| Training | Random seeds | 42, 43, 44 |
| Training | Optimizer | AdamW |
| Training | Learning rate | $10^{-3}$ |
| Training | Weight decay | $10^{-4}$ |
| Training | Gradient clipping | Maximum norm 5.0 |
| Training | Objective | Query-wise cross-entropy |
| Training | Training score | phenotype-gene evidence + gated graph score |
| Training | Initial adaptive gate value | 0.05 |
| Training | Negatives per query | 1,024 |
| Training | Positive-support masking probability | 0.20 |
| Training | Rescue-focused query fraction | 0.50 |
| Training | Effective query batch size | 32, via gradient accumulation |
| Training | Numerical precision | BF16 mixed precision |
| Training | Maximum epochs | 100 |
| Training | Early-stopping patience | 15 epochs |
| Training | Checkpoint selection | Highest validation Recall@50; MRR tie-break |

`47ab30a`, version `1.2.3-2-g47ab30a`, using the January 2021 H2GKB snapshot and its default skewness-based weighting.

##### S3.2 CADA

CADA learns node2vec embeddings on an HPO–gene knowledge graph augmented with clinical case associations and prioritises genes through link prediction. The evaluated configuration combines phenotype embeddings using an unweighted mean. CADA was run as version `0.0.1`, commit `56e42ec`, using its published unweighted G100 model and default parameters. The bundled HPO ontology was release 2019-09-06.

##### S3.3 GADO

GADO predicts HPO–gene associations from gene co-regulation estimated across large-scale RNA-seq data and combines phenotype-specific prediction scores. Its spiked prediction matrix retains known phenotype–gene associations. GADO was run as version `1.0.4` in PROCESS-then-Prioritise mode using its distributed resource bundle. The bundled HPO ontology was release 2020-08-11.

##### S3.4 Input and output handling

All methods received the observed HPO terms for each case. GADO and CADA discard terms absent from their bundled ontologies; where this left a case without a usable ranking, it was retained as a miss under the main-text evaluation policy. Cases whose causal gene was absent from the returned ranking were also retained as misses.

Ranked outputs were harmonised to a common gene-symbol namespace using the repository's mapping, and duplicate entries were removed. Rankings were evaluated in their returned order after this processing. Gene symbols were harmonized using the HGNC complete-set mapping distributed with the repository ("external/benchmark_tools/hgnc_complete_set.txt"), resolving unambiguous previous and alias symbols to current symbols. Duplicate entries were removed while preserving their first occurrence in the ranking.

All evaluated methods incorporate curated phenotype–gene knowledge, although through different mechanisms. PGRanker's distinction is therefore how it integrates that evidence: direct phenotype support initialises patient-specific gene representations, while an attention-pooled representation of the indexed HPO profile conditions scoring through gene–query interaction features.

#### S4 Additional results

##### S4.1 Phenotype degradation detailed results

Table S2 contains the detailed results for all four methods over all four phenotype degradation regimes applies.

##### S4.2 Mixed rare diseases benchmark per-case results

Table S3 contains the full per-case performance of all four methods on the 21-case mixed rare diseases benchmark.

Table S2: **Ranking performance under phenotype perturbations.** We report the area under the curve (AUC) for Recall@10 and mean reciprocal rank (MRR) under four perturbation regimes: removal, addition, substitution, and ancestor replacement. PGRanker achieves the highest AUC for both metrics across all four regimes.

| Regime | Metric | GADO | Phen2Gene | CADA | PGRanker (PGR) |
|---|---|---|---|---|---|
| Removal | Recall@10 AUC (%) | 6.03 | 10.18 | 21.29 | **25.86** |
| Addition | Recall@10 AUC (%) | 6.78 | 11.01 | 24.15 | **27.63** |
| Substitution | Recall@10 AUC (%) | 5.61 | 7.08 | 15.82 | **19.30** |
| Ancestor replacement | Recall@10 AUC (%) | 2.24 | 7.07 | 5.05 | **14.91** |
| Removal | MRR AUC | 0.0230 | 0.0444 | 0.1167 | **0.1752** |
| Addition | MRR AUC | 0.0237 | 0.0440 | 0.1350 | **0.1843** |
| Substitution | MRR AUC | 0.0210 | 0.0306 | 0.0882 | **0.1253** |
| Ancestor replacement | MRR AUC | 0.0095 | 0.0358 | 0.0303 | **0.0865** |

Table S3: Case-level causal-gene ranks and reciprocal ranks (RR) for all four methods on the 21-case mixed rare-disease cohort. The causal gene and number of HPO terms supplied as input are shown for each case. PhenGeneRanker results are averaged over seeds 42, 43, and 44, with ranks rounded to the nearest integer and RR to four decimal places. Boldface denotes the highest RR and lowest rank per case, determined before rounding. A dash indicates that the causal gene was absent from the returned ranking, assigned RR = 0.

| | | | PGRanker | | Phen2Gene | | CADA | | GADO | |
|---|---|---|---|---|---|---|---|---|---|---|
| Case ID | Causal gene | $n_{\text{HPO}}$ | RR | Rank | RR | Rank | RR | Rank | RR | Rank |
| 01 | VPS41 | 2 | **0.0049** | **204** | 0.0000 | – | 0.0000 | – | 0.0008 | 1301 |
| 02 | ROR2 | 4 | 0.0127 | 81 | **0.0323** | **31** | 0.0088 | 113 | 0.0033 | 301 |
| 03 | ACAN | 3 | 0.0021 | 499 | **0.0044** | **226** | 0.0012 | 851 | 0.0020 | 498 |
| 04 | TMEM67 | 3 | 0.0313 | 32 | 0.0071 | 141 | **0.0417** | **24** | 0.0088 | 113 |
| 05 | CHST3 | 3 | 0.0066 | 155 | 0.0003 | 3215 | **0.0093** | **108** | 0.0039 | 256 |
| 06 | NEK1 | 2 | **1.0000** | **1** | 0.0061 | 163 | 0.0076 | 131 | 0.0029 | 340 |
| 07 | TCTN3 | 3 | 0.0117 | 86 | 0.0071 | 140 | **0.1667** | **6** | 0.0035 | 285 |
| 08 | MESP2 | 2 | 0.0017 | 587 | **0.0089** | **112** | 0.0014 | 709 | 0.0014 | 718 |
| 09 | SLC12A1 | 3 | **0.0469** | **22** | 0.0133 | 75 | 0.0385 | 26 | 0.0208 | 48 |
| 10 | GAD1 | 2 | 0.0009 | 1080 | 0.0010 | 1006 | 0.0010 | 970 | **0.0016** | **607** |
| 11 | PPP1R21 | 3 | **0.1500** | **7** | 0.0001 | 14321 | 0.0000 | – | 0.0002 | 4156 |
| 12 | MAB21L1 | 4 | **0.0632** | **16** | 0.0006 | 1671 | 0.0075 | 134 | 0.0016 | 621 |
| 13 | TNFSF11 | 2 | 0.0011 | 877 | **0.0172** | **58** | 0.0007 | 1448 | 0.0021 | 483 |
| 14 | ECEL1 | 3 | **0.0372** | **33** | 0.0040 | 252 | 0.0116 | 86 | 0.0051 | 196 |
| 15 | POC1A | 1 | **0.0009** | **1185** | 0.0001 | 7578 | 0.0004 | 2831 | 0.0005 | 1866 |
| 16 | CCDC82 | 1 | **0.0003** | **3546** | 0.0000 | – | 0.0000 | – | 0.0001 | 7348 |
| 17 | TJP2 | 2 | **0.0942** | 11 | 0.0909 | **11** | 0.0286 | 35 | 0.0047 | 212 |
| 18 | ASPM | 1 | 0.0009 | 1064 | 0.0011 | 872 | **0.0286** | **35** | 0.0208 | 48 |
| 19 | MID2 | 1 | **0.0117** | **151** | 0.0001 | 6685 | 0.0016 | 613 | 0.0024 | 423 |
| 20 | TMEM67 | 3 | 0.0313 | 32 | 0.0071 | 141 | **0.0417** | **24** | 0.0088 | 113 |
| 21 | PSD | 1 | **0.0003** | **3417** | 0.0001 | 10907 | 0.0000 | – | 0.0001 | 8877 |

### References


[1] Mengge Zhao, James M Havrilla, Li Fang, Ying Chen, Jacqueline Peng, Cong Liu, Chao Wu, Mahdi Sarmady, Pablo Botas, Julián Isla, Gholson J Lyon, Chunhua Weng, and Kai Wang.

Phen2gene: rapid phenotype-driven gene prioritization for rare diseases. *NAR Genomics and Bioinformatics*, 2(2), May 2020.

[2] Jason A. O'Rawe, Yiyang Wu, Max J. Dörfel, Alan F. Rope, P.Y. Billie Au, Jillian S. Parboosingh, Sungjin Moon, Maria Kousi, Konstantina Kosma, Christopher S. Smith, Maria Tzetis, Jane L. Schuette, Robert B. Hufnagel, Carlos E. Prada, Francisco Martinez, Carmen Orellana, Jonathan Crain, Alfonso Caro-Llopis, Silvestre Oltra, Sandra Monfort, Laura T. Jiménez-Barrón, Jeffrey Swensen, Sara Ellingwood, Rosemarie Smith, Han Fang, Sandra Ospina, Sander Stegmann, Nicolette Den Hollander, David Mittelman, Gareth Highnam, Reid Robison, Edward Yang, Laurence Faivre, Agathe Roubertie, Jean-Baptiste Rivière, Kristin G. Monaghan, Kai Wang, Erica E. Davis, Nicholas Katsanis, Vera M. Kalscheuer, Edith H. Wang, Kay Metcalfe, Tjitske Kleefstra, A. Micheil Innes, Sophia Kitsiou-Tzeli, Monica Rosello, Catherine E. Keegan, and Gholson J. Lyon. Taf1 variants are associated with dysmorphic features, intellectual disability, and neurological manifestations. *The American Journal of Human Genetics*, 97(6):922–932, Dec 2015.

[3] Jung Hoon Son, Gangcai Xie, Chi Yuan, Lyudmila Ena, Ziran Li, Andrew Goldstein, Lulin Huang, Liwei Wang, Feichen Shen, Hongfang Liu, Karla Mehl, Emily E. Groopman, Maddalena Marasa, Krzysztof Kiryluk, Ali G. Gharavi, Wendy K. Chung, George Hripcsak, Carol Friedman, Chunhua Weng, and Kai Wang. Deep phenotyping on electronic health records facilitates genetic diagnosis by clinical exomes. *The American Journal of Human Genetics*, 103(1):58–73, Jul 2018.

[4] Chao Wu, Batsal Devkota, Perry Evans, Xiaonan Zhao, Samuel W. Baker, Rojeen Niazi, Kajia Cao, Michael A. Gonzalez, Pushkala Jayaraman, Laura K. Conlin, Bryan L. Krock, Matthew A. Deardorff, Nancy B. Spinner, Ian D. Krantz, Avni B. Santani, Ahmad N. Abou Tayoun, and Mahdi Sarmady. Rapid and accurate interpretation of clinical exomes using phenoxome: a computational phenotype-driven approach. *European Journal of Human Genetics*, 27(4):612–620, Jan 2019.

[5] Cong Liu, Fabricio Sampaio Peres Kury, Ziran Li, Casey Ta, Kai Wang, and Chunhua Weng. Doc2hpo: a web application for efficient and accurate hpo concept curation. *Nucleic Acids Research*, 47(W1):W566–W570, May 2019.

[6] Atteeq U. Rehman, Maryam Najafi, Marios Kambouris, Lihadh Al Gazali, Periklis Makrythanasis, Abolfazl Rad, Reza Maroofian, Anna Rajab, Zornitza Stark, Jill V. Hunter, Zeineb Bakey, Mari J. Tokita, Weimin He, Francesco Vetrini, Andrea Petersen, Federico A. Santoni, Hanan Hamamy, Kaman Wu, Fatma Al Jasmi, Martin Helmstädter, Sebastian J. Arnold, Fan Xia, Christopher Richmond, Pengfei Liu, Ehsan Ghayoor Karimiani, GholamReza Karami Madani, Sebastian Lunke, Hatem El-Shanti, Christine M. Eng, Stylianos E. Antonarakis, Jozef Hertecant, Magdalena Walkiewicz, Yaping Yang, and Miriam Schmidts. Biallelic loss of function variants in PPP1R21 cause a neurodevelopmental syndrome with impaired endocytic function. *Human Mutation*, 40(3):267–280, Mar 2019.

[7] Abolfazl Rad, Umut Altunoglu, Rebecca Miller, Reza Maroofian, Kiely N. James, Ahmet Okay Çağlayan, Maryam Najafi, Valentina Stanley, Rose-Mary Boustany, Gözde Yeşil, Afsaneh Sahebzamani, Gülhan Ercan-Sencicek, Kolsoum Saeidi, Kaman Wu, Peter Bauer, Zeineb Bakey, Joseph G. Gleeson, Natalie Hauser, Murat Gunel, Hulya Kayserili, and Miriam Schmidts. MAB21L1 loss of function causes a syndromic neurodevelopmental disorder with distinctive cerebellar, ocular, craniofacial and genital features (COFG syndrome). *Journal of Medical Genetics*, 56(5):332–339, May 2019.

[8] Leslie E. Sanderson, Kristina Lanko, Maysoon Alsagob, Rawan Almass, Nada Al-Ahmadi, Maryam Najafi, Mohammad A. Al-Muhaizea, Hamad Alzaidan, Hesham AlDhalaan, Elena Perenthaler, Herma C. van der Linde, Anita Nikoncuk, Nikolas A. Kühn, Dinu Antony, Tarek Mustafa Owaidah, Salmo Raskin, Luana Gabriela Dalla Rosa Vieira, Romulo Mombach, Najmeh Ahangari, Tainá Regina Damaceno Silveira, Najim Ameziane, Arndt Rolfs, Aljohara Alharbi, Raghda M. Sabbagh, Khalid AlAhmadi, Bashayer Alawam, Hazem Ghebeh, Aljouhra AlHargan, Anoud A. Albader, Faisal S. Binhumaid, Ewa Goljan, Dorota Monies, Osama M. Mustafa, Mazhor Aldosary, Albandary AlBakheet, Banan Alyounes, Faten Almutairi, Ali Al-Odaib, Durdane Bekar Aksoy, A. Nazli Basak, Robin Palvadeau, Daniah Trabzuni, Jill A. Rosenfeld, Ehsan Ghayoor Karimiani, Brian F. Meyer, Bedri Karakas, Futwan Al-Mohanna, Stefan T. Arold, Dilek Colak, Reza Maroofian, Henry Houlden, Aida M. Bertoli-Avella, Miriam Schmidts, Tahsin Stefan Barakat, Tjakko J. van Ham, and Namik Kaya. Bi-allelic variants in HOPS complex subunit VPS41 cause cerebellar ataxia and abnormal membrane trafficking. *Brain*, 144(3):769–780, Mar 2021.

[9] Chengyao Peng, Simon Dieck, Alexander Schmid, Ashar Ahmad, Alexej Knaus, Maren Wenzel, Laura Mehnert, Birgit Zirn, Tobias Haack, Stephan Ossowski, Matias Wagner, Theresa Brunet, Nadja Ehmke, Magdalena Danyel, Stanislav Rosnev, Tom Kamphans, Guy Nadav, Nicole Fleischer, Holger Fröhlich, and Peter Krawitz. Cada: phenotype-driven gene prioritization based on a case-enriched knowledge graph. *NAR Genomics and Bioinformatics*, 3(3), June 2021.

[10] Patrick Deelen, Sipko van Dam, Johanna C. Herkert, Juha M. Karjalainen, Harm Brugge, Kristin M. Abbott, Cleo C. van Diemen, Paul A. van der Zwaag, Erica H. Gerkes, Evelien Zonneveld-Huijssoon, Jelkje J. Boer-Bergsma, Pytrik Folkertsma, Tessa Gillett, K. Joeri van der Velde, Roan Kanninga, Peter C. van den Akker, Sabrina Z. Jan, Edgar T. Hoorntje, Wouter P. te Rijdt, Yvonne J. Vos, Jan D. H. Jongbloed, Conny M. A. van Ravenswaaij-Arts, Richard Sinke, Birgit Sikkema-Raddatz, Wilhelmina S. Kerstjens-Frederikse, Morris A. Swertz, and Lude Franke. Improving the diagnostic yield of exome- sequencing by predicting gene–phenotype associations using large-scale gene expression analysis. *Nature Communications*, 10(1), June 2019.
